# Molecular architecture and platelet-activating properties of small immune complexes assembled on intact heparin and their possible involvement in heparin-induced thrombocytopenia

**DOI:** 10.1101/2023.02.11.528150

**Authors:** Yang Yang, Yi Du, Daniil Ivanov, Chendi Niu, Rumi Clare, James W. Smith, Ishac Nazy, Igor A. Kaltashov

## Abstract

Heparin-induced thrombocytopenia (HIT) is an adverse reaction to heparin leading to a reduction in circulating platelets with an increased risk of thrombosis. It is precipitated by polymerized immune complexes consisting of pathogenic antibodies that recognize a small chemokine platelet factor 4 (PF4) bound to heparin, which trigger platelet activation and a hypercoagulable state. Characterization of these immune complexes is extremely challenging due to the enormous structural heterogeneity of such macromolecular assemblies and their constituents (especially heparin). We use native mass spectrometry to characterize small immune complexes formed by PF4, heparin and monoclonal HIT-specific antibodies. Up to three PF4 tetramers can be assembled on a heparin chain, consistent with the results of molecular modeling studies showing facile polyanion wrapping along the polycationic belt on the PF4 surface. Although these assemblies can accommodate a maximum of only two antibodies, the resulting immune complexes are capable of platelet activation despite their modest size. Taken together, these studies provide further insight into molecular mechanisms of HIT and other immune disorders where anti-PF4 antibodies play a central role.

## Introduction

Platelet factor 4 (PF4) is a chemokine that is secreted from activated platelets^1^ and involved in a variety of physiological processes, ranging from coagulation and tissue repair to innate immune response.^2^ It has gained particular notoriety due to its role in triggering a range of (auto)immune hematologic pathologies, the first of which - heparin-induced thrombocytopenia (HIT) – was initially reported sixty-five years ago.^3^ More recently it was implicated in thrombotic complications accompanying rare but potentially fatal side effects of adenoviral -vector COVID-19 vaccines, presently known as the vaccine-induced immune thrombotic thrombocytopenia (VITT).^4^ A common feature of these disorders is formation of large immune complexes, in which multiple PF4 tetramers polymerize anti-PF4 antibodies. This leads to clustering of FcγRIIa receptors on the platelet surface, thereby initiating cross-phosphorylation of the Immunoreceptor Tyrosine-based Activation Motifs (ITAMs) within their cytosolic parts,^5^ which is the first step in the platelet activation cascade.^6^ While the sequence of events leading to the onset of HIT has been known for some time, the actual occurrence of this pathology among heparin patients (including those that produce anti-PF4 antibodies) appears to be random and impossible to predict (as is the occurrence of VITT among vaccinated individuals).

The structure of the PF4-based immune complexes must be one of the major determinants of the occurrence of HIT and the progression of this pathology, but its detailed characterization at the molecular level is extremely challenging due to both macro-heterogeneity of such assemblies and micro-heterogeneity of their constituents. The macro-heterogeneity refers to the broad distribution of sizes exhibited by both immune complexes and their precursors – PF4/heparin assemblies.^7^ The latter are usually classified based on their elution behavior in size exclusion chromatography (SEC) as either small or ultra-large complexes (SCs and ULCs, respectively). SCs are considered precursors to ULCs, and elucidation of the architecture of the SCs (involving only a single heparin chain) is important not only for understanding their specific role in HIT pathogenesis, but also for building a molecular model of ULCs. Both SCs and ULCs exhibit a range of sizes and stoichiometries, although there is evidence suggesting that equimolar PF4:heparin stoichiometry is optimal vis-à-vis formation of ULCs, which are more pathogenic.^8^

The micro-heterogeneity refers to the structural diversity within the constituents of both SCs and ULCs, which is mostly due to the variation of heparin chain length and its sulfation and acetylation patterns,^9,10^ although some variability within the circulating PF4 has also been noted.^11^ Because of the extensive heterogeneity exhibited by the PF4/heparin complexes, their initial characterization relied on relatively low-resolution methods of structural analysis, such as size-exclusion chromatography (SEC).^8,12,13^

Meaningful utilization of high-resolution methods capable of providing detailed structural information is also possible, but requires both the size and the heterogeneity of the relevant macromolecular assemblies to be reduced dramatically, *e*.*g*. to enable their crystallization and recording of interpretable diffraction patterns. This was recently accomplished by Cai et al.,^14^ who not only solved the crystal structure of recombinant PF4 complexed to a small (pentasaccharide) synthetic heparinoid, fondaparinux, but was also able to obtain an atomic-level structure of the fondaparinux-bound PF4 associated with a Fab segment of a monoclonal anti-PF4/heparin antibody (KKO)^15^ that mimics the behavior of human HIT antibodies. One important feature of the PF4/fondaparinux complexes revealed in that study is the allosteric change within the PF4 tetramers induced by the short polyanion binding, which exposes a neo-epitope on the protein surface and enables its recognition by KKO (and, by extension, pathogenic wild-type HIT antibodies).^14^ However, because of the small size of the complex and the homogeneity of its constituents (unlike heparin, fondaparinux is chemically defined, and its length is only 1/10 of the average heparin chain), the structure of the pathogenic PF4/heparin complexes could not be determined directly, but instead was inferred by extrapolating the features of the relatively small and homogeneous PF4/fondaparinux assemblies to a larger scale.^14^ One particularly intriguing conclusion was that PF4 tetramer binding to the heparin chain “*imparts a local linearized structure*” within the latter, thereby enhancing binding of the second PF4 molecule.^14^ It was suggested that progression of this process should lead tо formation of a large antigenic complex, in which PF4 tetramers cluster around a semi-rigid heparin chain.^14^ This scenario differs from the earlier models of PF4/heparin complexes, in which the anionic polysaccharide is wrapped around the tetramer along its equatorial belt where the basic amino acids are localized.^16^ The conformational bias towards the linearized chains upon heparin/PF4 association envisioned by Cai and co-workers^14^ has important ramifications, as it would allow high-density protein tethering to a single polyanion chain, which would enable effective platelet activation by immune complexes assembled on a single heparin compared to the classical models^16^ (which imply a much lower number of PF4 tetramers bound to a single heparin chain – and, therefore, a lower number of anti-PF4/heparin antibodies that can associate with such complexes). However, characterization of the molecular architecture of clinically relevant PF4/heparin assemblies and HIT-related immune complexes based upon them remained outside of the reach of experimental biophysics.

In the past two decades mass spectrometry (MS) proved to be a powerful tool for characterization of non-covalent biopolymer assemblies; however, its utility in the field of protein-glycosaminoglycan (GAG) interactions remains confined to systems with limited size and modest structural heterogeneity.^17-19^ The progress made in the field of MS-based analysis of highly heterogeneous systems, and particularly incorporation of ion chemistry in the MS workflow^20^ (such as the limited charge reduction technique^21^) resulted in a dramatic expansion of the scope of macromolecules and their assemblies for which meaningful information can be obtained. In this work we employ native MS supplemented with limited charge reduction to study association of PF4 with intact unfractionated heparin (UFH), revealing a range of binding stoichiometries within SCs. Although the largest assembly within this set corresponds to only three PF4 tetramers accommodated by a single heparin chain (consistent with the classical model of PF4/heparin interaction^16^ and confirmed by molecular modeling), the corresponding immune complexes may incorporate up to two antibody molecules, as revealed by native MS. Importantly, the results of platelet activation assays show that such immune complexes are capable of triggering platelet activation, and are a surrogate marker for the HIT disease. This poses an intriguing question of whether the minimal size of the FcγRIIa clusters on the platelet surface required for its activation is indeed two, or the initial immune complex incorporating only two antibody molecules serves only as a seed for a large FcγRIIa cluster formation on the platelet surface, a process facilitated by the dimensionality reduction (from 3D in circulation to 2D on the cell surface). The picture of the SC molecular organization emerging from this work also provides important insights vis-à-vis architecture of ULCs, the MDa-size PF4/heparin assemblies that are commonly considered to be the critical element in triggering HIT.

## Results

### PF4/heparin association: polyanion chain length as the major determinant of the SC architecture and evidence supporting the “wrap-around” interaction model

Close approximation of at least two PF4 tetramers is a key to formation of an immune complex capable of activating the platelet,^13^ although it remains unclear if there is a minimal heparin chain length that enables such protein clustering. Previously we have shown that short heparin fragments (at least up to a decasaccharide level, dp10) are not sufficient for PF4 bridging (instead, multiple heparinoids associate with a single PF4 tetramer).^22^ Similar behavior is exhibited by significantly longer chains, such as dp20 (the length that corresponds to an average size of the low-molecular weight heparin products, LMWH^23^), as can be seen in **Figure 1A**. Incubation of dp20 with PF4 at an approximately equimolar ratio gives rise to an abundant ionic signal corresponding to a PF4·(dp20)_2_ complex, although the 1:1 protein/heparinoid complexes are also observed (**Figure 1A, top**). Mass measurements of the most abundant species in both distributions indicates that average mass of dp20 species incorporated in the PF4·(dp20)_2_ complex is slightly higher compared to that within the PF4·dp20 complex (the average mass of the protein-bound dp20 increases by up to 170 Da), indicating the preference for more extensively sulfated species within the former. Higher concentration of dp20 in the protein/heparinoid mixture results in a near-complete elimination of the PF4·dp20 complexes, and a signal shift for the PF4·(dp20)_2_ species towards higher *m/z* values (**Figure 1A, bottom**). This mass increase (0.25-0.3 kDa) is indicative of the increased extent of sulfation of the protein-bound heparinoid chains (three to four “extra” sulfate groups per each heparinoid chain), consistent with the notion of the high charge density being the most important factor vis-à-vis heparin binding preferences to polycationic proteins.^24^ We also note the absence of the putative complexes formed by bridging of two (or more) PF4 tetramers by a single dp20 chain. The surprisingly limited range of stoichiometries exhibited by the PF4/dp20 complexes is corroborated by the molecular modeling results, which provide clear indication that the positive charge basin around the PF4 tetramer can readily accommodate up to two dp20 chains (**Figure 1B**). The favorable ratio of the heparin chain persistence length to the protein dimensions allows the dp20 chain to maximize the electrostatic interactions by circumscribing the PF4 surface.

**Figure 1.**
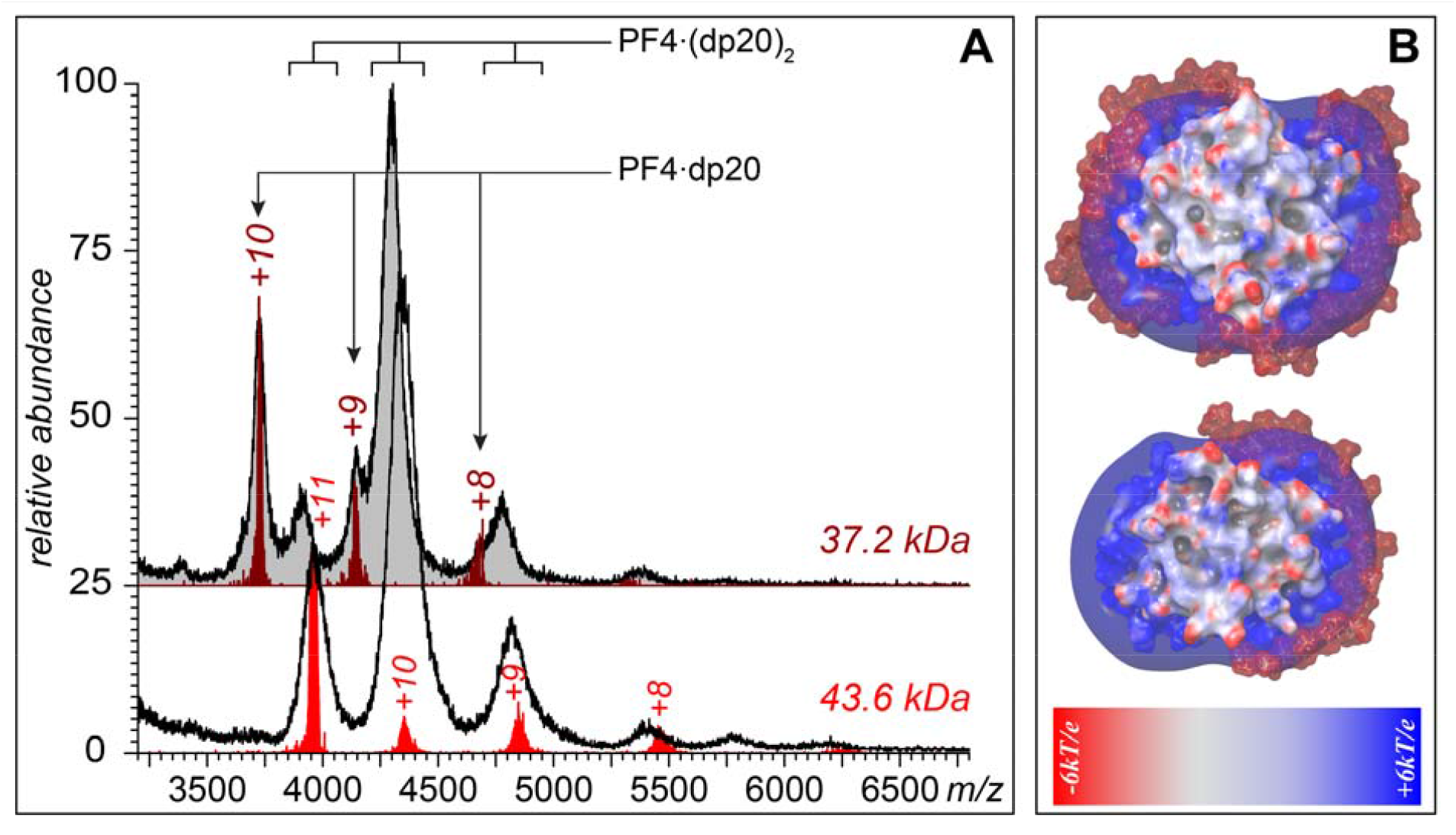
**A**: Native MS of PF4/dp20 mixtures (***top***: 0.27 and 0.028 mg/mL, respectively; ***bottom***: 0.27 and 0.045 mg/mL). The maroon and red charge ladders were obtained by limited charge reduction of ionic populations at *m/z* 3,723 (top) and 3,953 (bottom). The two masses calculated based on these charge ladders correspond to 1:1 (37.2 kDa) and 1:2 (43.6 kDa) protein:heparinoid stoichiometries. **B**: Molecular modeling of PF4/dp20 interactions showing 1:1 and 1:2 protein/heparinoid complexes. The iso-potential surface (6*k*T/*e*) represents the positive belt around the protein, while the heparin chain is shown using a space-fill model (without the corresponding iso-potential surface).

Binding of multiple (up to three) PF4 tetramers to a single polyanion chain becomes evident only when unfractionated heparin (UFH) is used as an assembly scaffold. Consistent with a previous study,^8^ the SEC chromatogram of a PF4/UFH mixture (**Figure 2A**) displays two distinct peaks representing ULC (elution window 6-7 min) and SC (9.5-12 min). Fractionation of the chromatographic peak representing SCs enables isolation of PF4/heparin assemblies of varying dimensions (**Figure 2A**), although the stoichiometries of the putative (PF4)_*n*_·(heparin)_m_ complexes comprising each subfraction cannot be assigned based on the elution time alone.

**Figure 2.**
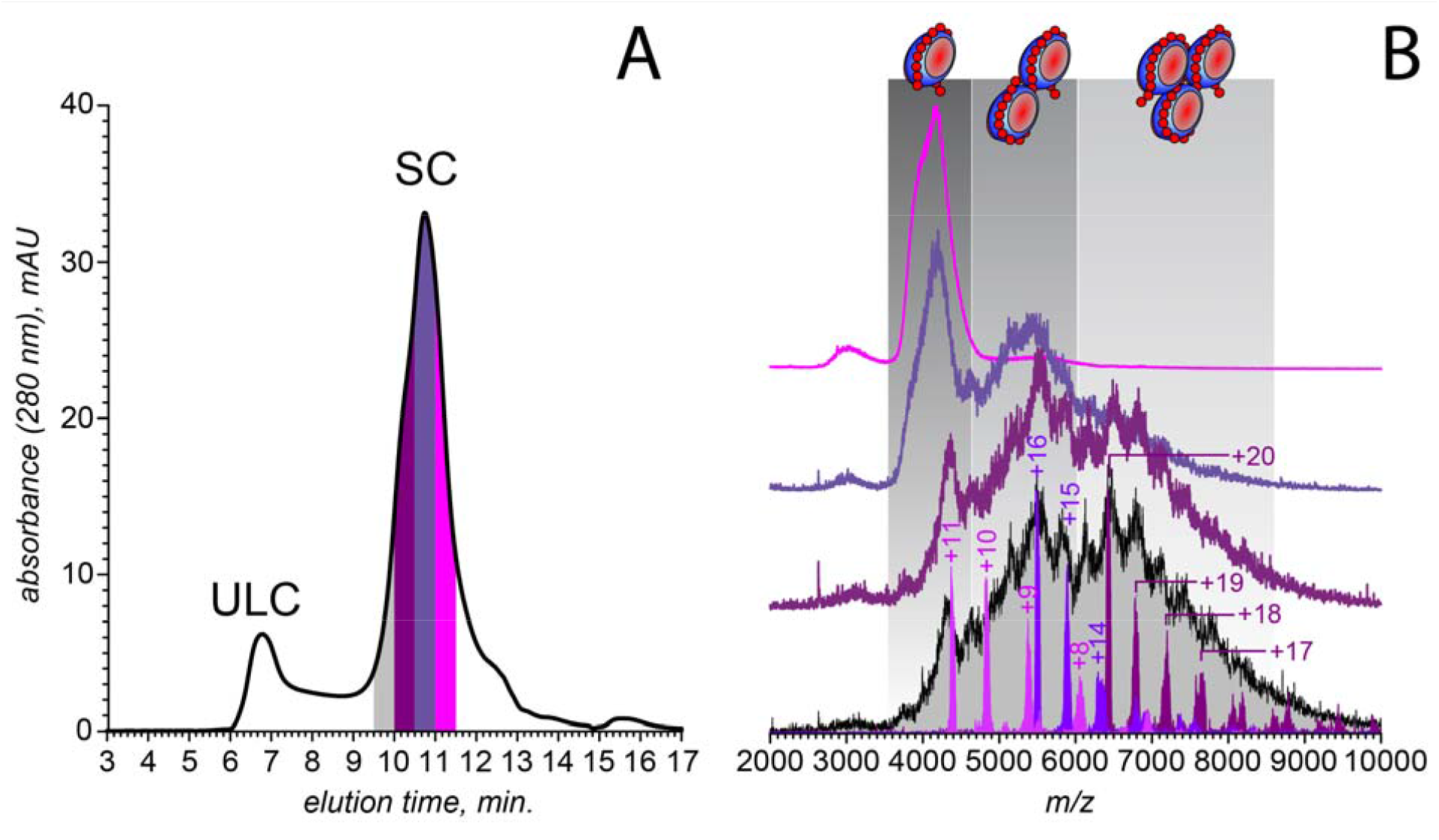
**A**: SEC of PF4 incubated with UFH (1.6 mg/mL and 0.67 mg/mL, respectively). **B**: native MS

Native MS analysis of the latest-eluting subfraction (collection window 11.0-11.5 min) yields a mass spectrum that contains an abundant ionic signal within a relatively narrow *m/z* range (3,500-4,500), but does not display any resolved charge states. The three other subfractions give rise to even more convoluted ionic signals, which gradually shift towards higher *m/z* values. Overall, three discernable features are observed in the mass spectra of the four SC subfractions (confined to *m/z* regions 3,500-4,500; 4,500-6,000 and 6,000-8,000), indicating the presence of three distinct species in solution that differ from each other in size. The abundance of the lowest-*m/z* species gradually declines across the collected subtractions, while the highest-*m/z* species can only be observed within the two early-eluting SC subfractions. Despite the convoluted appearance of these mass spectra, the ionic masses can be readily determined using limited charge reduction. For example, application of this technique to interpret the spectral feature localized within the *m/z* region 3,500-4,500 reveals its identity as 1:1 PF4/heparin complexes. These measurements are illustrated in **Figure 2B** with a charge ladder produced by exposing the ionic population isolated at *m/z* 4,383 to radical anions, which results in facile loss of up to four elementary charges. This charge ladder allows the mass of these ionic species to be calculated as 48.5 kDa, suggesting that the average mass of the heparin chain incorporated in these complexes is 16.8 kDa. Moving the precursor ion selection window to the *m/z* region 4,500-6,000 (the second distinct spectral feature in **Figure 2B**) followed by limited charge reduction of the selected ions also gives rise to a well-defined charge ladder. The mass range of the corresponding ionic species is 82-88 kDa, indicative of the presence of two PF4 tetramers within the complexes, and the mass of the heparin component being 20-26 kDa (which corresponds to chains comprised of 66-86 fully sulfated saccharides). Lastly, the masses of the ionic species giving rise to the third spectral feature are determined by selecting the precursor ions at *m/z* 6422. Limited charge reduction of these ions yields a 122-129 kDa mass range, indicative of the presence of three PF4 tetramers within the complex, and the mass of the heparin component falling within the 29-36 kDa range (96-120 saccharide unit-long chains). Although these numbers appear high compared to the average size of UFH (15 kDa, or 50 monosaccharide units^25,26^), size profiling of UHF using intact-mass MS supplemented with limited charge reduction reveals a broad distribution with the upper mass limit exceeding 40 kDa (see ***Supplementary Material***).

The measured masses of SCs with various heparin/PF4 stoichiometries indicate that accommodation of a single PF4 tetramer by a heparin chain requires segments consisting of 30-40 monosaccharide units. Although this protein placement density is lower compared to that predicted by extrapolating the crystal structure of the PF4/fondaparinux complex,^14^ molecular modeling of PF4 interactions with long heparin chains (dp 40, dp70 and dp100) indicates that the polyanion possesses sufficient flexibility to wrap around the PF4 tetramers, thereby maximizing the electrostatic interactions (**Figure 3**). The protein molecules in these structures are not only almost completely circumscribed with heparin chains, but also brought into proximity with each other such that the entire polyanion chain interacts with the positive charge basins on the proteins’ surfaces.

**Figure 3.**
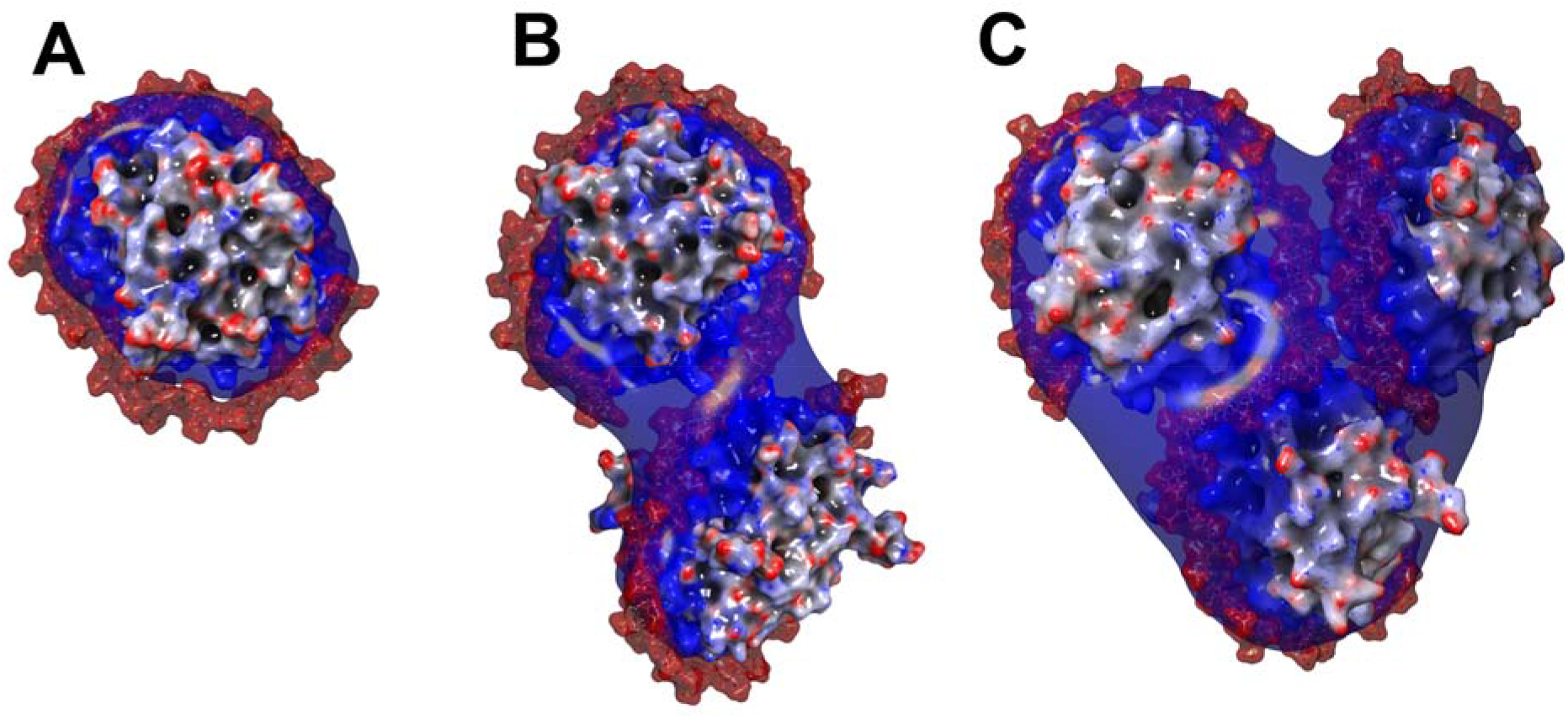
Molecular modeling of dp40/PF4 (**A**), dp70/PF4 (**B**) and dp100/PF4 (**C**) interactions yielding 1:1, 1:2 and 1:3 heparin/PF4 complexes, respectively. The iso-potential surfaces (**A**: 5.5 kT/e and **B, C**: 7.5*k*T/*e*) represent the positive belts around the protein, while the heparin chain is shown using a space-fill model (without a corresponding iso-potential surface). Refer to ***Supplementary Material*** for details of the molecular modeling work.

### Immune complexes formed by SC association with model HIT antibodies

The mass spectrum of PF4/dp20 complexes incubated with a nearly-equimolar amount of the KKO antibodies displays an abundant ionic signal above *m/z* 6000 (**Figure 4**) that can be deconvoluted to yield three components: the-lowest mass species (152.5±1 kDa) corresponds to the antigen-190 free KKO molecules, while two others have masses consistent with KKO·PF4·(dp20)_2_ species (195.5±1 kDa), and the antigen-saturated antibody KKO·[PF4·(dp20)_2_]_2_ and [dp20·PF4]·KKO·[PF4·(dp20)_2_] (237±5 kDa). No immune complexes contain more than a single antibody molecule, consistent with the notion of a single neo-epitope generated on the PF4 surface following its association with heparin.^14^ Au contraire, incubation of the highest-MW sub-fraction of SCs (having the highest PF4 load – up to three protein tetramers per single heparin chain, see **Figure 2**) with a nearly-equimolar amount of KKO gives rise to abundant ionic signal extending well beyond *m/z* 10,000 (**Figure 5**). While some spectral features can be readily recognized as the free antibody (*m/z* 6,000-7,500) and unconsumed SCs (*m/z* 4,000-6,000), assignment of the abundant ionic signal at m/z 7,500-9,000 cannot be made using native MS alone due to the lack of distinct features corresponding to individual charge states. Limited charge reduction allows the mass of the most abundant ions within this spectral region to be determined as 240 kDa (the red trace in **Figure 5**). This value falls within the expected mass range for the 1:2:1 heparin/PF4/KKO complexes (see ***Supplementary Material***). Interestingly, the most abundant signal (populating the high-*m/z* region of the mass spectrum) shows clearly discernable features representing individual charge states, which allows the ionic mass to be calculated as 424±2.7 kDa, a value that corresponds to a 1:3:2 heparin/PF4/KKO complex. No complexes of 1:3:3 heparin/PF4/KKO stoichiometry were detected even when the antibody was present in molar excess (up to 33%).

**Figure 4.**
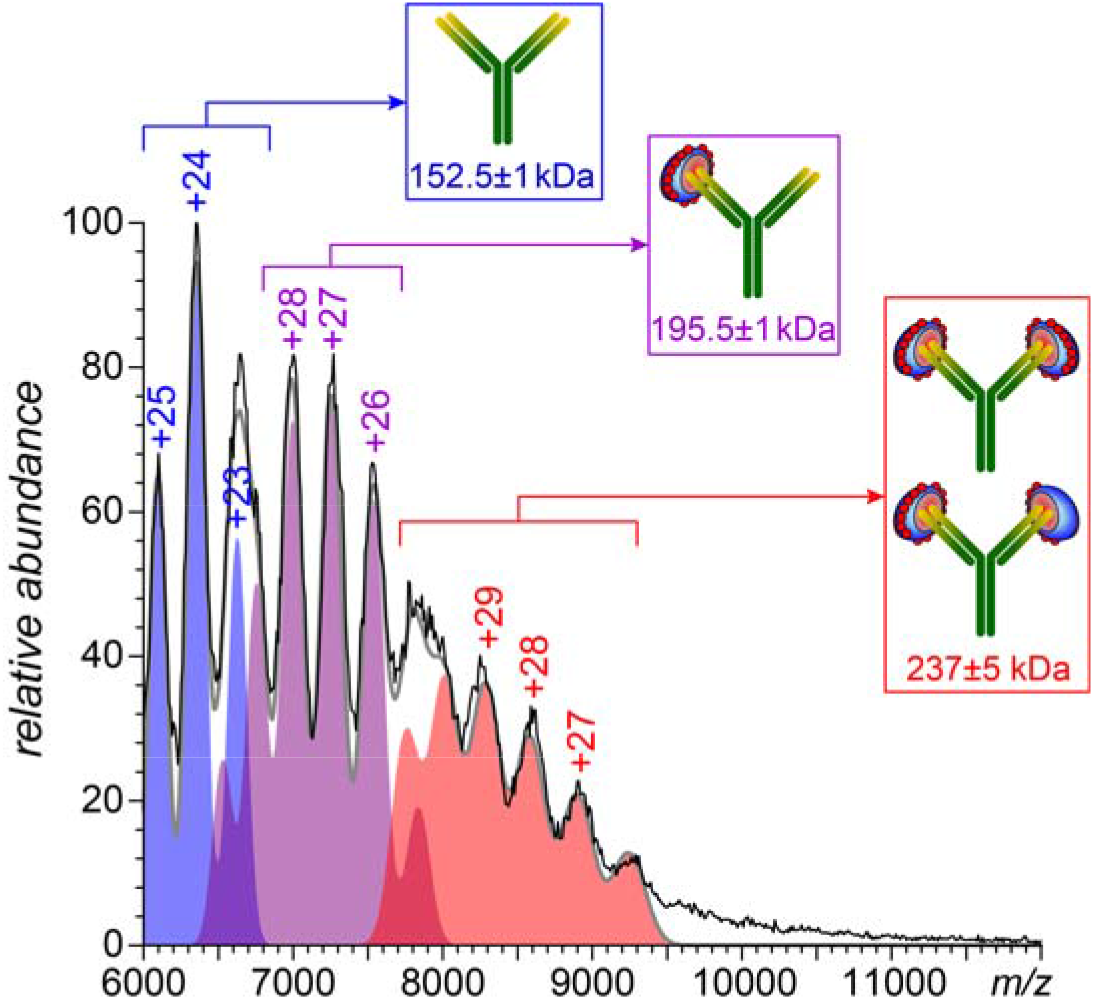
Native MS of PF4/dp20 complexes incubated with KKO (0.28 mg/mL and 0.06 mg/mL, respectively). The color-filled curves show contributions of three distinct species to the overall ionic signal (the sum of these contributions is shown with a gray trace).

**Figure 5.**
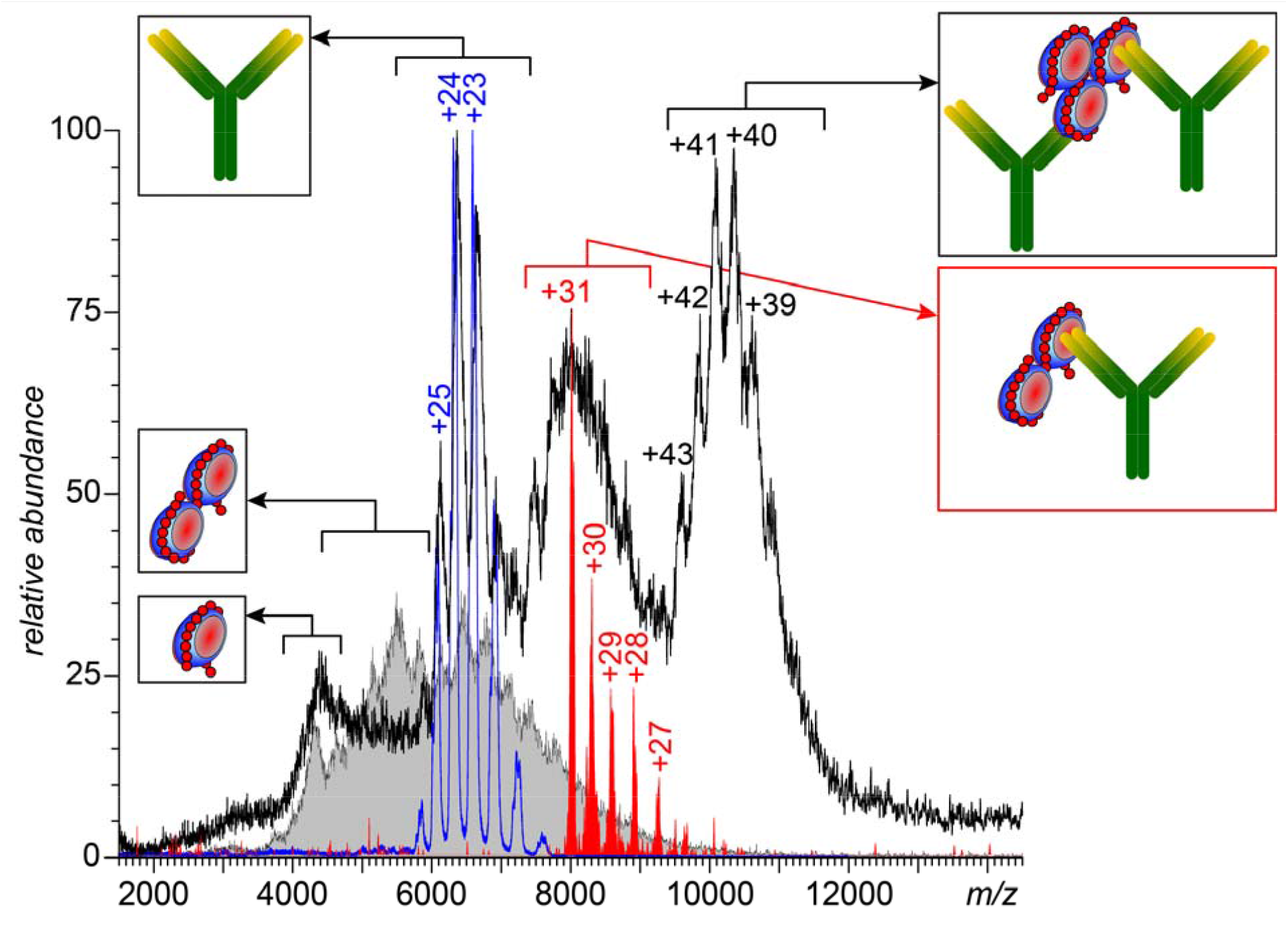
Native MS of an immune complex formed by KKO and the highest-MW sub-fraction of SCs. The blue and gray traces represent reference spectra of KKO and the highest-MW subfraction of SCs, respectively. The red trace represents the charge ladder produced by limited charge reduction of the ionic population selected at *m/z* 8,000.

### Biological activity of the immune complexes built on SC templates

Small immune complexes prepared by incubation of KKO with a UFH/PF4 mixture were fractionated to produce components with different antibody content (**Figure 6**). Consistent with the native MS data (*vide supra*), SEC chromatogram of the incubated mixture features two partially resolved peaks indicating the presence of either two or one KKO molecules in the immune complexes. The corresponding fractions (designated as A and B, respectively) were tested using serotonin release assay (SRA) with platelets collected from two donors. Even though borderline activity was detected for the crude sample prior to its fractionation (see the blue bars in **Figure 6**), it is the fraction containing immune complexes with two antibody molecules per complex (fraction A) that showed the highest platelet-activating potential regardless of the nature of PF4 used to prepare the initial UFH/PF4 mixture (*i*.*e*., both the recombinant PF4 and the platelet-derived PF4). While the SRA carried out with the platelets collected from another donor showed lower overall activity, the highest levels of serotonin release were once again registered for fraction A (see ***Supplementary Material***).

**Figure 6.**
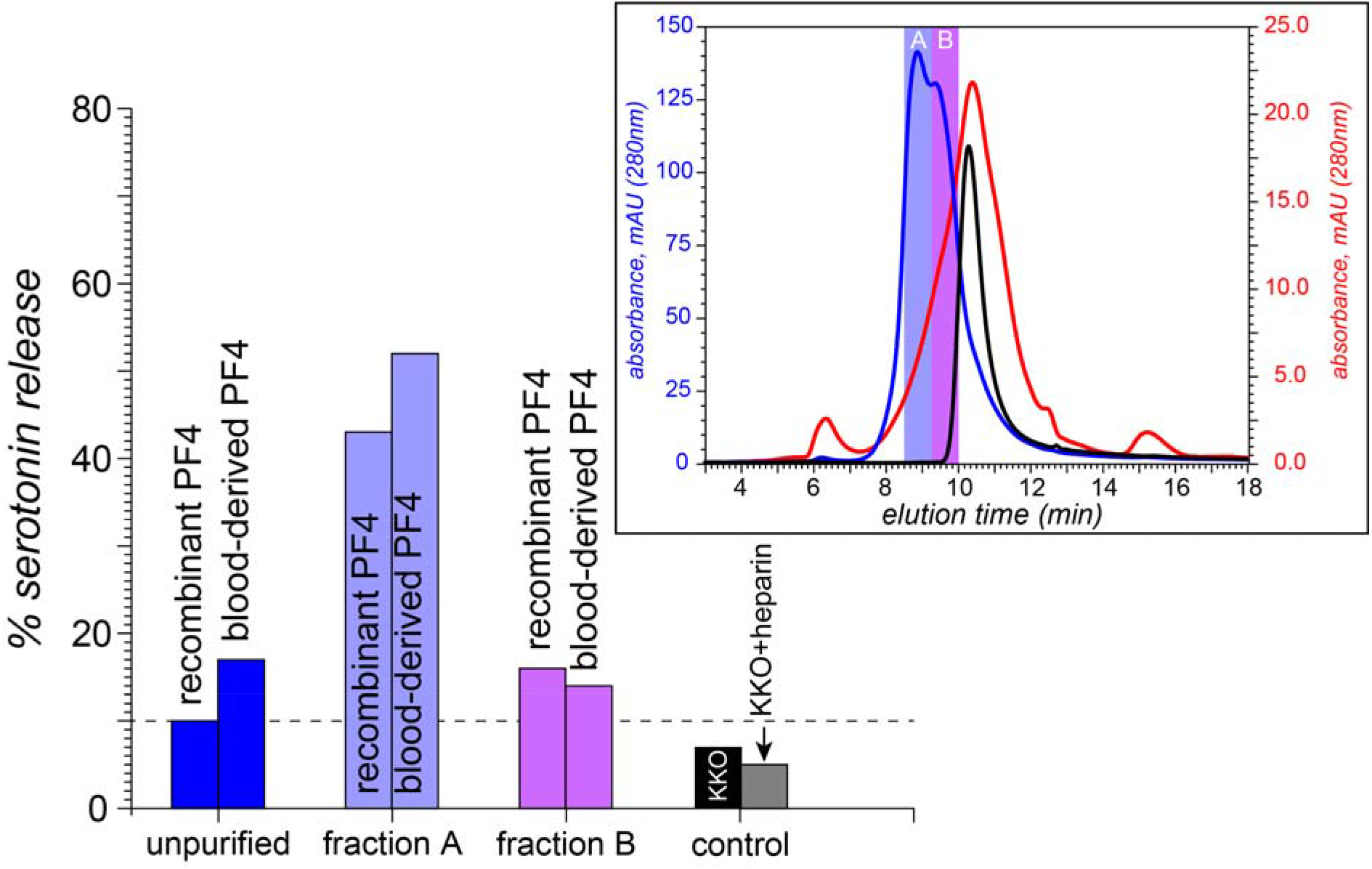
SRA data showing the platelet activation by KKO incubated with a UFH/PF4 mixture without fractionation of the immune complexes (blue bars), and after isolating fractions containing two and one KKO molecules per immune complex (fractions A and B represented with the violet and purple bars, respectively); platelet activation is shown as the percent release of ^14^C-serotonin. The left and right bars in each group represent the data acquired with recombinant and blood-derived forms of PF4, respectively. The control measurements are shown with the black and gray bars (KKO alone and KKO/heparin mixture, respectively). The chromatogram in the inset (blue trace) shows the elution windows during which the two fractions (A and B) were collected; the reference chromatograms of the

## Discussion

Current understanding of the GAG/protein interactions invokes the notion of a multiplicity of mechanisms ranging from highly specific associations that are defined by surface complementarity (analogous to protein/protein binding) to non-specific binding that appears to lack any selectivity.^27,28^ Electrostatic interactions are the most important forces driving protein association with polyanionic GAGs. The binding affinity is frequently determined by how well localization of the sulfate groups within the GAG chains matches the distribution of basic residues on the protein surface,^27^ a concept which is frequently referred to as a “sulfation code.”^29,30^ Heparin is the most densely charged GAG (on average, it carries three sulfate groups per disaccharide unit in addition to a single carboxylate). The high charge density and its relatively even distribution across the heparin chain (unlike heparan sulfate, where short highly sulfated segments are spaced by relatively long low charge-density regions^9,31^) provides for an ideal match to the positive charge distribution across the PF4 surface (which forms an equatorial belt around the protein tetramer,^16,32^ giving rise to nearly symmetric toroidally-shaped isopotential surfaces). This naturally invites a suggestion that a sufficiently long heparin chain should envelope the protein molecule to maximize the enthalpic gains due to favorable electrostatic interactions.^16^ Even though heparin maintains extended/stretched conformations in solution,^33,34^ its chain flexibility is sufficient to allow the PF4 tetramer to be neatly circumscribed along its positive charge belt (*i*.*e*., heparin’s Kuhn segment length of 9±2 nm^35^ is notably lower compared to the PF4 tetramer diameter of 52±3 nm).

The favorable flexibility parameters of the heparin chains and their high charge density suggest that the “wrap-around” mode of PF4/heparin interactions is favored thermodynamically, which is also confirmed by the results of MD simulations (**Figure 3**). Although the MD simulations carried out in this study were not intended to explore the entire conformational space available to PF4/heparin complexes (explicit modeling remains problematic for GAG chains above decamers^28,36,37^), they certainly demonstrate that the “wrap-around” mode of association is feasible. At the same time, one must also consider a possibility of kinetic effects preventing formation of structures that maximize the electrostatics-driven enthalpic gains. For example, initial random polyanion/protein contacts may give rise to metastable structures in which the electrostatic interactions are not maximized, but the kinetic barriers separating them from the lower-energy minima are sufficiently high to guarantee their survival for long time periods. While such a scenario (a “Velcro effect”) appears feasible, the experimentally observed correlation between the heparin chain length and the number of PF4 tetramers it can accommodate provides a strong argument against it. Indeed, native MS indicates that each PF4 tetramer in heparin·(PF4)_n_ complexes recruits a segment consisting of 35-40 saccharide units, which is indeed the length required to circumscribe the entire protein along its positive charge belt (**Figure 3**). The thermodynamically favored polyanion collapse onto the polycationic surface of PF4 also explains the failure of the relatively short heparinoids to bridge two PF4 tetramers (unlike proteins with less extensive basins of the positive charge - such as neutrophil elastase - which can form dimers bridged by heparinoids as short as decasaccharides^38^). Our previous work^22^ demonstrated that such heparinoids (dp10) are incapable of PF4 bridging,^22^ and the results of the present study indicate that even heparin fragments that are twice as long (dp20) are not sufficiently large for PF4 bridging.

While the average length of UFH chains is ca. 50 saccharides, MS analysis assisted by limited charge reduction reveals a broad size distribution with the largest (but less abundant) heparin molecules exceeding 40 kDa, which corresponds to chains that are over 120 saccharide unit-long (see ***Supplementary Material***). This length is sufficient to allow complete wrapping of three PF4 tetramers by a single heparin chain, yielding heparin·(PF4)_3_ assemblies (the largest ones observed in MS measurements, **Figure 3**). The absence of the heparin/PF4 assemblies with higher protein load among SCs is fully consistent with the “wrap-around” model of the protein/polyanion interaction (higher binding stoichiometry would require heparin molecules that are sufficiently longer than those present in UFH).

The tight wrapping of the protein molecules by heparin chains within the SCs revealed by native MS and molecular modeling tools provides an explanation for another intriguing observation, namely the mismatch between the number of PF4 tetramers and antibodies (KKO) within the largest immune complexes built upon SCs. Indeed, the tight packing of the protein molecules within SCs results in steric hindrance that prevents association of more than a single antibody with PF4 on either side of the complex (**Figure 7**). In this case, KKO binding to the (PF4)_3_·heparin complex is possible only in the trans conformation, thereby limiting the antibody load of such complexes to two, as observed by native MS (**Figure 5**). Despite their modest antibody load, such complexes are capable of activating platelets as shown by the results of *in vitro* testing (**Figure 6**). This raises an intriguing question: is the close approximation of only two FcγRIIa receptors via binding to a single KKO_2_·(PF4)_3_·heparin complex sufficient for triggering platelet activation, or is the receptor pairing only a first step in assembling large FcγRIIa clusters, driven by polymerization of the platelet-bound KKO_2_·(PF4)_3_·heparin complexes on the cell surface (**Figure 8**). The ability of KKO to promote oligomerization of PF4/heparin complexes has been previously proposed,^39^ and even though the KKO-mediated polymerization of SCs in solution was not observed in our work, it is possible that it is accelerated on the platelet surface due to the dimensionality reduction (from 3D in solution to 2D on the platelet surface). Should this “on-the-membrane” polymerization of small immune complexes occur, it would culminate in the platelet activation triggered by FcγRIIa clustering and the ensuing phosphorylation cascade.^5,40^ The resulting platelet degranulation and release of PF4 will create a positive feed-back loop in HIT progression.^41,42^ Furthermore, there is a growing realization that in addition to platelets, pathogenesis of HIT-related thrombosis involves other blood cells,^43^ most of which constitutively express FcγRIIa^44^ and, therefore, can be activated by the heparin/PF4-templated immune complexes. Neutrophils are particularly important players in this respect due to their ability to release neutrophil extracellular traps (NETs)^45^ upon activation, which – due to the high content of long polyanions, such as DNA and polyphosphates – can act synergistically with heparin to provide additional scaffolds for assembling large PF4/polyanion complexes.

**Figure 7.**
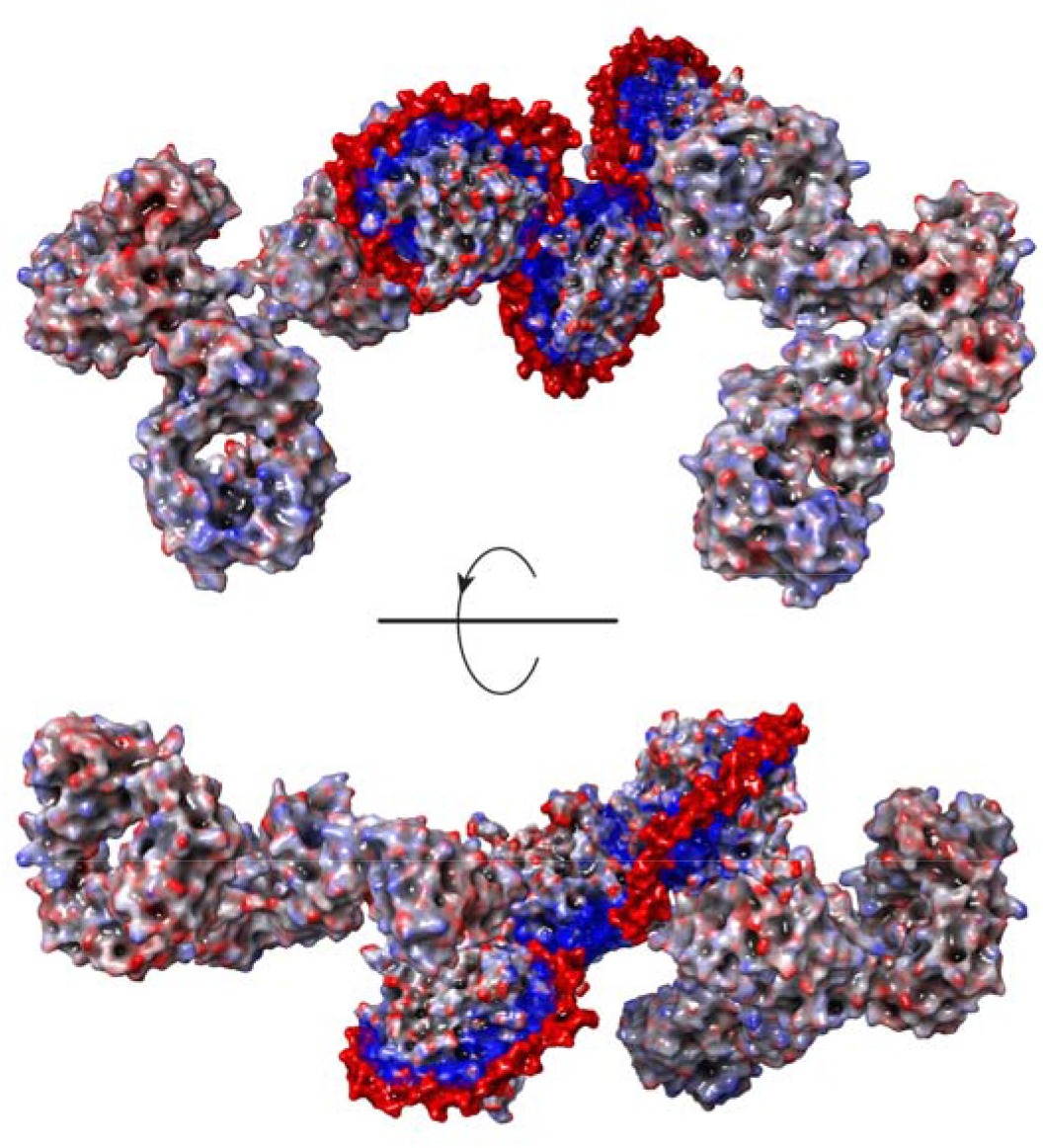
A representative conformation of the dp100·(PF4)_3_·KKO_2_ complex obtained by docking two KKO Fab segments to the dp100·(PF4)3 complex followed by a 500 ns MD simulation and energy minimization. Both Fab segments in the energy-minimized structure were extended to the full-length IgG molecules followed by another cycle of energy minimization.

**Figure 8.**
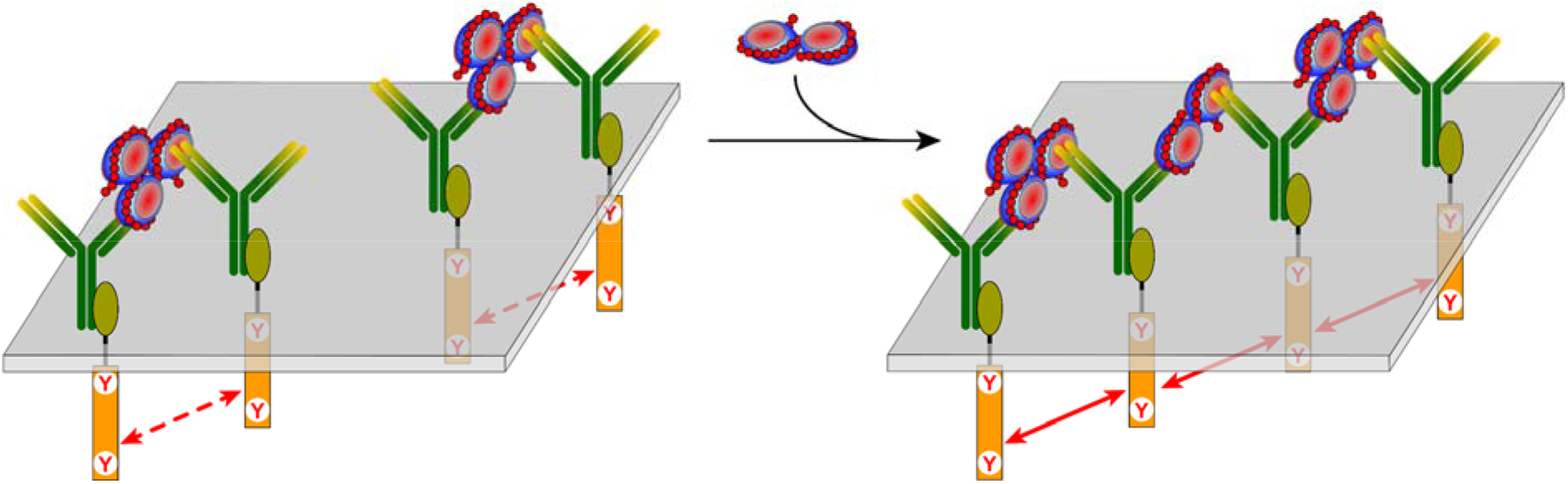
A schematic representation of two scenarios of platelet activation by immune complexes built upon SC/KKO association. ***Left***: KKO_2_·(PF4)_3_·heparin, the largest SC/KKO complexes produced in solution, bind to the FcγRIIa receptors on the platelet surface, which leads to pairing of the ITAM segments and enables their activation. ***Right***: the receptor-bound KKO_2_·(PF4)_3_·heparin complexes are cross-linked by SCs via the “vacant” Fab arms of the antibody molecules, giving rise to large FcγRIIa clusters on the platelet surface and close localization of multiple ITAM segments.

Our study has been focused primarily on the small PF4/heparin complexes, while the ULCs largely remained beyond its scope. However, the SC architecture revealed in this work provides important clues vis-à-vis the ULC structure and its assembly process. Since even the longest heparin chains cannot accommodate more than three PF4 tetramers, formation of the large complexes (exceeding 0.65 MDa by early estimates^8^ and averaging *ca*. 3 MDa according to our own estimates – see ***Supplementary Material***) must proceed through association of multiple heparin chains and PF4 tetramers. Since heparin chains maximize the electrostatic interactions with PF4 by circumscribing it, the heparin/PF4 complex polymerization process would proceed beyond the 1:3 stoichiometry (the observed SC limit) only if at every stage of the process there is a mismatch between the length of the unaccommodated heparin segment and the circumference of the PF4 tetramer being added to this nascent complex (see ***Supplementary Material***). Once these two dimensions become comparable, the growth of the complex stops, since neither GAG nor protein components of the complex will have any vacant high-charge density segments left to support further growth. This model is consistent with the notion of the ULC composition conforming to 1:1 stoichiometry, and our rough estimations of the ULC mass based on the model that correlates ionic *m/z* value and charge (*z*) in native MS^46^ yield ca. 3 MDa (***Supplementary Material***), suggesting that the number of PF4 tetramers in a single ULC particle exceeds 50. Even if a significant proportion of these molecule is inaccessible to anti-PF4 antibodies due to the steric hindrance, these particles should be able to accommodate a large number of antibodies and potentially lead to formation of large FcγRIIa clusters on the platelet surface. Nevertheless, the ability of the SC-based immune complexes to activate platelets (**Figure 8**) should not be overlooked. This mechanism does not appear to be unique to HIT, but is likely involved in other pathologies precipitated by platelet activation with antibody cross-linking by polyvalent antigens, such as VITT.^6^

## Conclusions

Combination of native MS with molecular modeling allows molecular architecture to be established for the PF4/heparin small complexes and the immune complexes templated on SCs. Heparin chains possess sufficient flexibility to wrap around polycationic PF4 molecules to maximize electrostatic attraction. This mode of interaction dictates the maximum size of SCs (up to three protein molecules bound to a heparin chain), as accommodation of a single PF4 tetramer within the complex requires a polyanionic segment consisting of 30-40 saccharide units. Despite the presence of up to three PF4 molecules within SCs, no more than two anti-PF4 antibodies associate with a single SC. Surprisingly, even this relatively low level of antibody loading is sufficient to trigger platelet activation. It remains to be seen whether bringing only two FcγRIIa receptors on the platelet surface into a close proximity with each other and the ensuing cross-phosphorylation of their ITAM elements activates the platelet by setting in motion the Syk pathway^47^ or further assembly of large receptor clusters on the cell surface is precipitated by polymerization of small immune complexes driven by the dimensionality reduction. Although the highly pathogenic ULCs largely remain beyond the scope of native MS at present, the structural properties of SCs revealed in this study provide important clues as to how these large, MDa-sized PF4/heparin assemblies are produced. The continuous improvements in native MS technology and expansion of its scope leave no doubt that the emerging model of ULC architecture will be tested in the near future, allowing the mechanisms of HIT and related thrombotic disorders to be thoroughly studied at the molecular level.

## Methods

### Materials

Both UFH and dp20 were purchased from Galen Laboratory Supplies (North Haven, CT). The recombinant form of human PF4 was expressed and purified using a protocol described elsewhere,^48^ and blood-derived human PF4 was obtained from Hematologic Technologies (now Prolytix, Essex Junction, VT). The KKO mAb was obtained using KKO hybridoma clone 31.2.57.A7 cell line (ATCC Manassas, VA). These cells were first thawed in HCELL-100 hybridoma serum free media (Wisent Bioproducts, Saint Jean-Baptiste, QC, Canada) supplemented with 200 mM L-glutamine (Thermo Fisher Scientific, Waltham, MA) and 10 000 U/mL penicillin + 10 000 µg/mL streptomycin (Thermo Fisher Scientific) then transferred to T-25 vented tissue culture flasks (Sarstedt, Nümbrecht,Germany) and incubated at 37°C and 5% CO_2_ for 3 days. KKO cells were then diluted 1/10 and passaged every 3-5 days. Passages 3, 4 and 5 were overgrown for an additional 10-14 days after which the supernatant was collected and frozen at -80°C. KKO was then purified using Protein A Sepharose beads (Thermo Fisher Scientific) packed in an Econo-chromatography column (BioRad, Hercules, CA) and washed 3 times with phosphate buffered saline (PBS). Thawed KKO supernatant was then run through the Protein A column and after the beads were washed with PBS, KKO antibody was eluted with 0.1 M Glycine pH 2.8 in 1 mL aliquots and immediately neutralized with 100 µL of 1.5M Tris pH 8.8. Next, the optical density (OD) at 280 nm was measured using a spectrophotometer (Eppendorf, Hamburg, Germany) and individual fractions with KKO were assessed for purity using SDS-PAGE and pooled. Finally, the concentration of the pooled fractions was measured and KKO binding to biotinylated PF4 was assessed using an in-house enzyme immunoassay. The quality of all biological products used in this study was verified using MS or LC/MS analysis. PF4/heparin and PF4/dp20 complexes were prepared by incubation of the protein/GAG mixtures in 150 mM ammonium acetate at 37 °C for 30 min using mixing stoichiometries as described in the text. SEC fractionation of SCs was carried out using a Sepax Nanofilm SEC-500 (Sepax, Newark, DE) on an HP1200 (Agilent Technologies, Santa Clara, CA) chromatograph. All protein and heparin solutions for MS analyses were prepared in 150 mM ammonium acetate, pH adjusted to 6.9.

### Mass Spectrometry

All native MS measurements were carried out using a Synapt G2Si (Waters Corp., Milford, MA) hybrid quadrupole/time-of-flight mass spectrometer equipped with a nanospray ionization source. The following set of parameters was used in the ESI interface region to ensure the survival of non-covalent complexes: capillary voltage, 1.4 kV; sampling cone voltage, 80 V; source offset, 80 V; trap CE, 4 V; trap DC bias, 3 V; and transfer CE, 0 V. Isolation of ionic populations in the trap cell for subsequent limited charge reduction measurements was performed by setting the quadrupole LM resolution values in the range of 4.0–4.5. Charge reduction of the selected polycationic ions was triggered by introducing 1,3-dicyanobenzene anions after setting the trap wave height to 0.2 V and optimizing the discharge current.

### In vitro platelet activation studies

Two sets of PF4/heparin complexes were prepared and tested for their ability to activate platelets using serotonin release assays (SRA),^49^ one containing the recombinant form of human PF4, and the other containing platelet-derived PF4. In each case PF4 was incubated with UHF (final concentrations 0.126 and 0.285 mg/mL, respectively) at 37 °C for 30 min in 1X PBS buffer, followed by addition of KKO (final concentration 3.4 mg/mL). The resulting immune complexes were fractionated on SEC, and the two most abundant fractions were tested twice using platelets from two different donors. The control tests were carried out using 1.95-2.24 mg/mL KKO without PF4/heparin complexes in the media.

### Molecular modeling

Molecular modeling of GAG/protein complexes was carried out with Desmond (Schrödinger Release 2022-2, Schrödinger LLC, New York, NY) using the OPLS4 force field.^50^ Sodium and chloride ions were added to neutralize any charges and to obtain 150 mM salt concentration. The PF4 tetramer and the KKO Fab segment extracted from PDB 4R9Y were used as the initial structures in the modeling studies. PF4 tetramer was further modified by adding the missing acidic N-termini. Fully sulfated heparin chains of different lengths were extracted from PDB 3IRJ. The dp70·(PF4)_2_model was built from energy-minimized dp70 chain and two individual PF4; the distance between dp70 and PF4 tetramers of the starting frame were selected to exceed 50 Å to avoid any artificial interactions. The structure was further simulated (MD) for 300 ns. The dp100·(PF4)_3_ structure was built using energy-minimized dp100 chain as a template and two PF4 tetramers positioned close to the GAG chain ends. This structure was simulated for 60 ns followed by addition of the third PF4 tetramer. This structure was further simulated (MD) for additional 300 ns.

## Authors Contributions

YY prepared the samples, carried out MS measurements and interpreted the data; YD prepared the samples for biological testing and carried out molecular modeling work; DI carried out MS measurements and interpreted the data; CN carried out MS measurements; RC expressed the recombinant forms of PF4 and KKO; JS carried out the platelet activation measurements and interpreted the data; IN designed the biological testing of the small immune complexes and provided oversight for the recombinant protein production; IK designed the study, provided oversight of the experimental work (MS measurements), interpreted the data and drafted the manuscript. All co-authors have participated in editing the manuscript and gave their consent to its final (submitted) version.

## Acknowledgements

This work was supported by a grant R01 GM112666 from the National Institutes of Health.

## Supplementary Material

**Figure S1.**
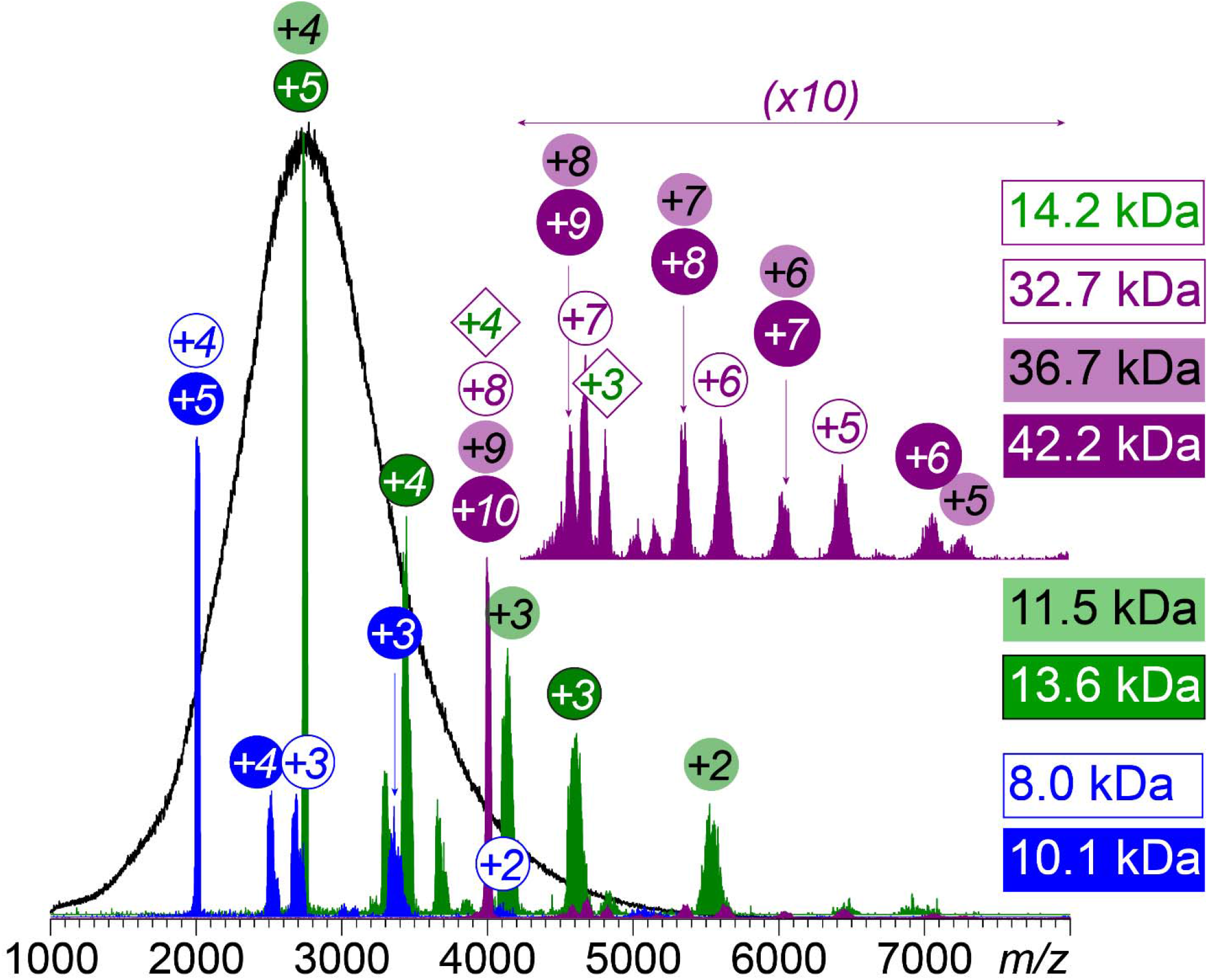
Native MS of UFH (0.25 mg/mL in 150 mM ammonium acetate) and mass assignment for representative ionic populations based on limited charge reduction measurements.

**Table S1.**
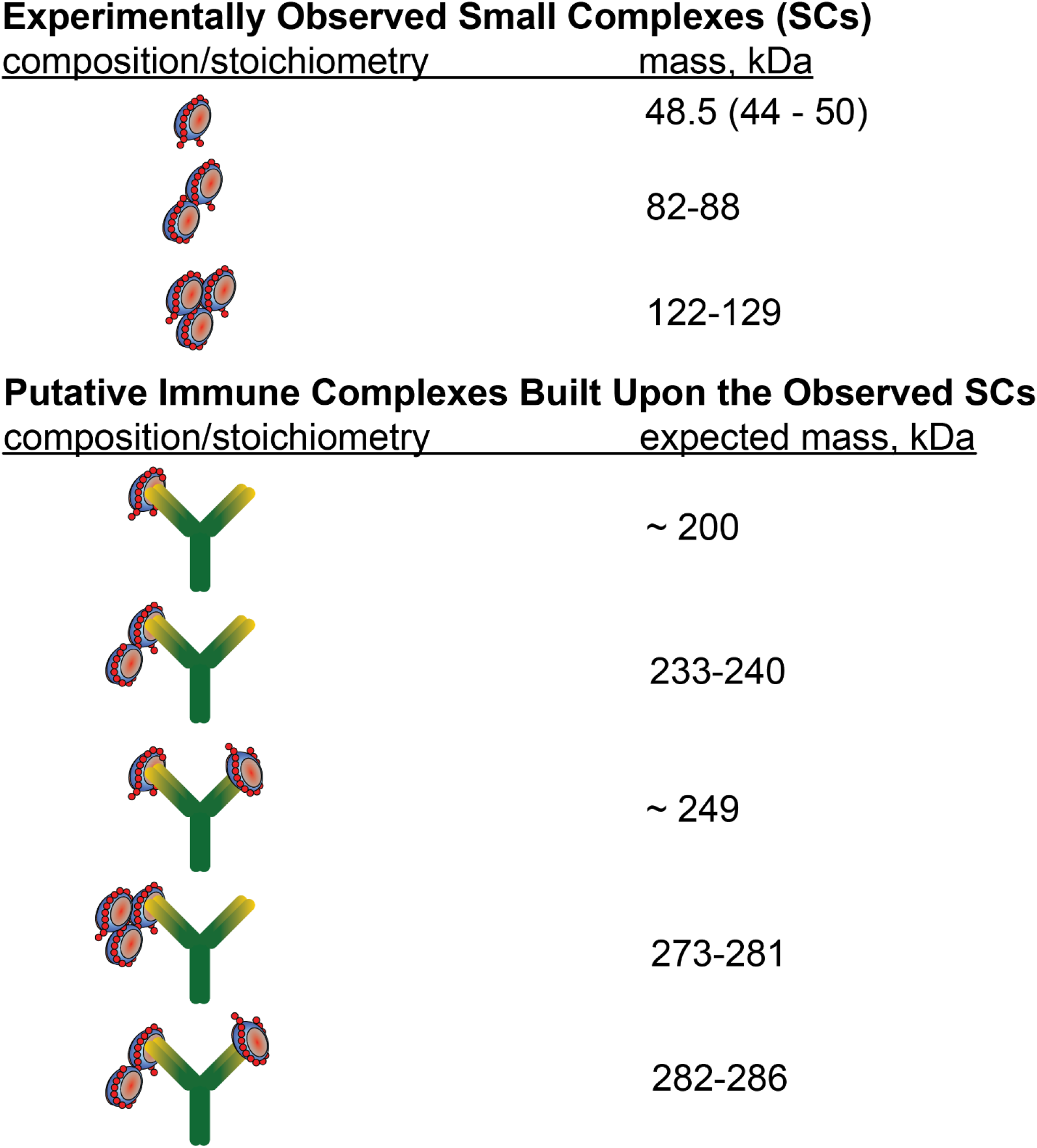
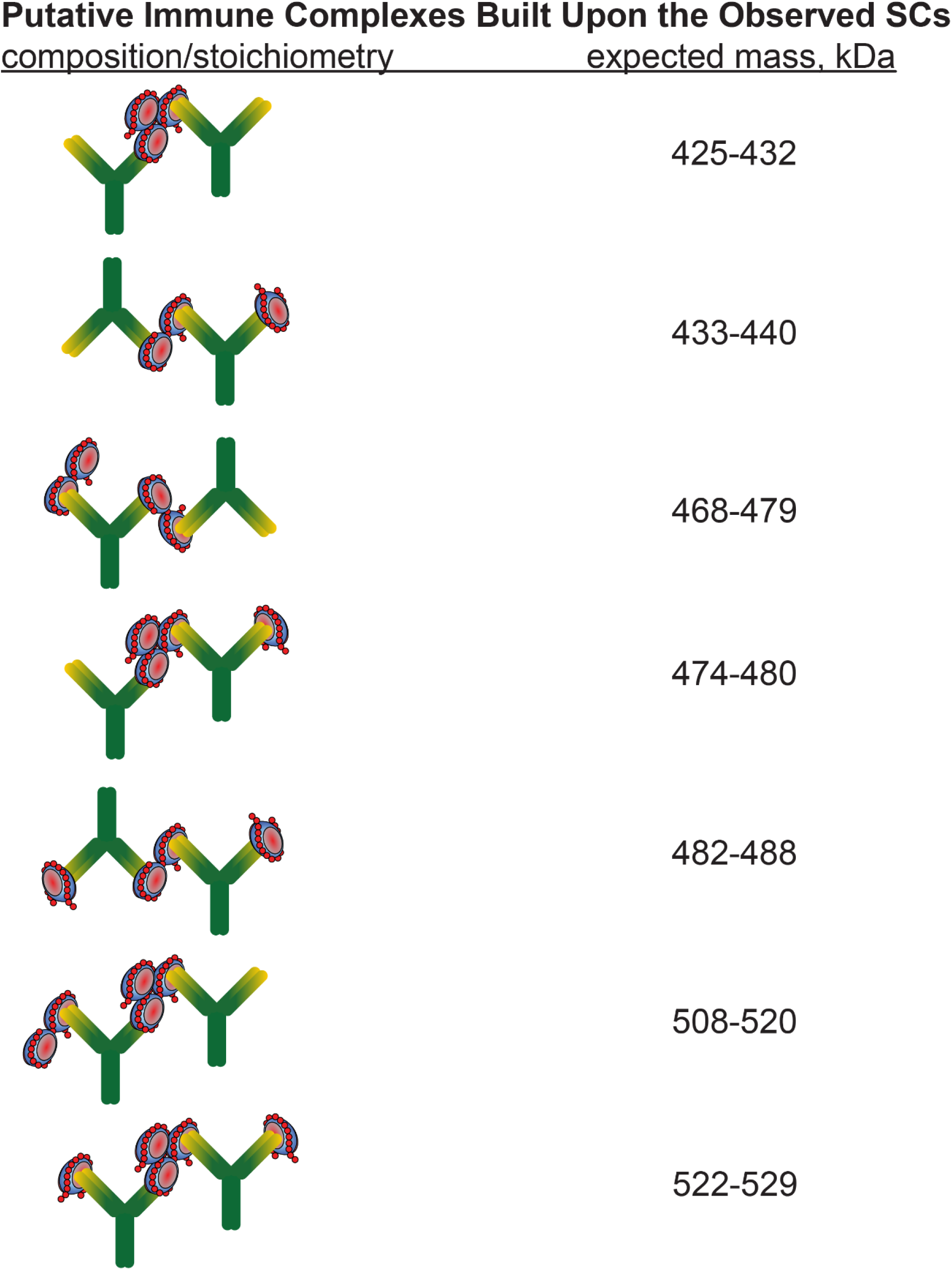
Expected mass ranges for UFH/PF4/KKO complexes with different stoichiometries.

**Figure S2.**
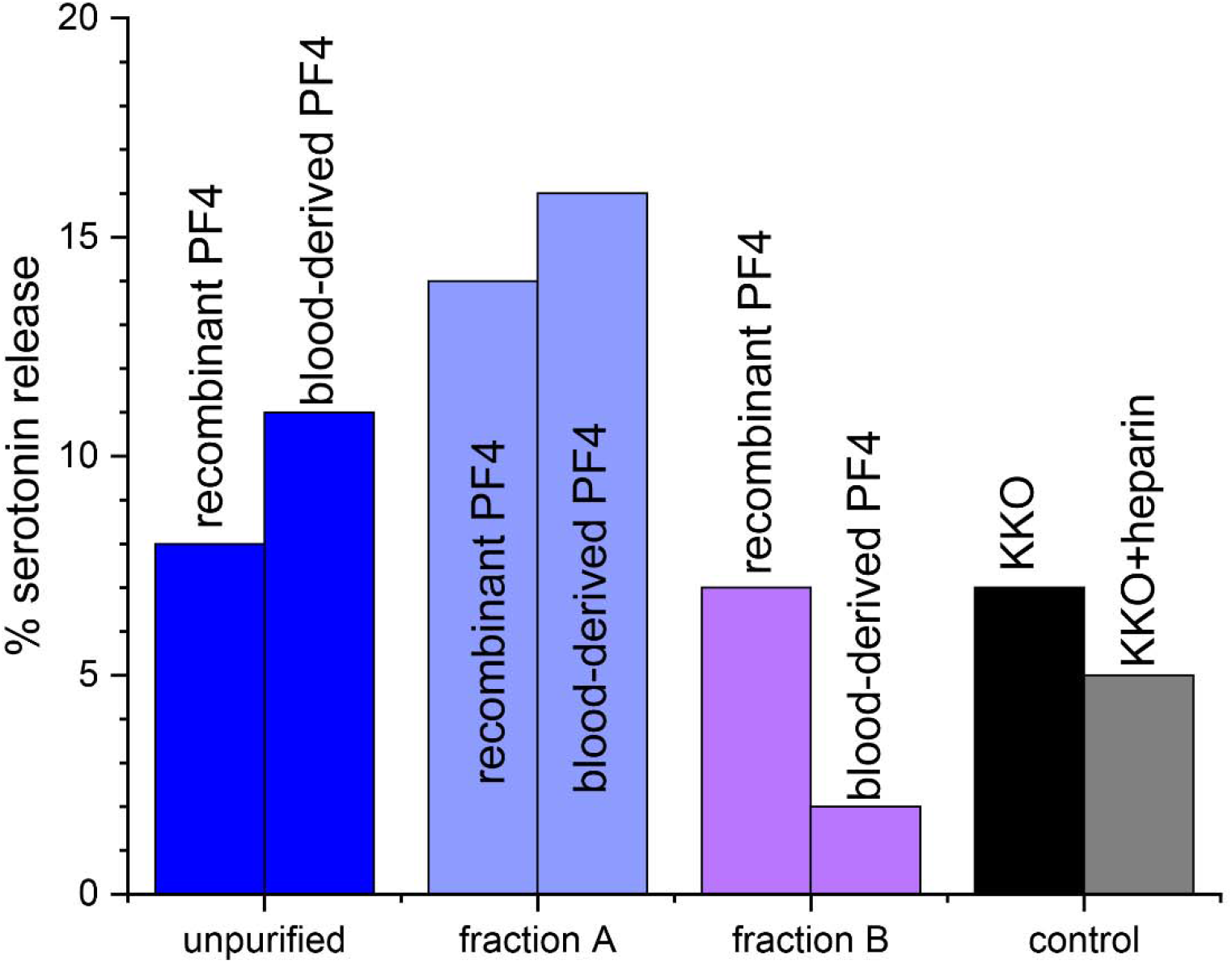
SRA data showing the platelet activation by KKO incubated with a UFH/PF4 mixture without fractionation of the immune complexes (blue bars), and after isolating fractions containing two and one KKO molecules per immune complex (fractions A and B represented with the violet and purple bars, respectively); platelet activation is shown as the percent release of ^14^C-serotonin. The left and right bars in each group represent the data acquired with recombinant and blood-derived forms of PF4, respectively. The control measurements are shown with the black and gray bars (KKO alone and KKO/heparin mixture, respectively). Platelets used in the SRA assay were collected from Donor 2.

**Figure S3.**
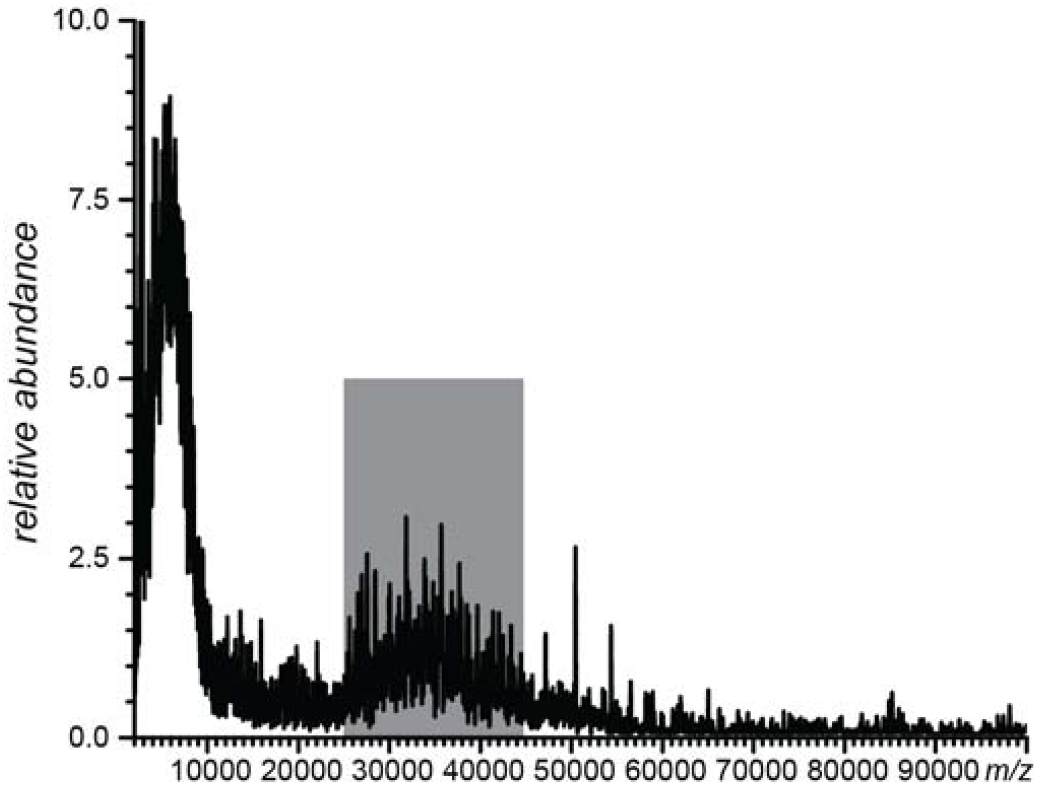
Native MS of ULCs generated in the UFH/PF4 mixture (0.67 mg/mL and 1.6 mg/mL, respectively). The unresolved ionic signal in the mass spectrum (*m/z* range 25,000-45,000) was used to estimate the average ULC mass (3.0±1.5 MDa) following the approach described elsewhere.^1^

Briefly, the mass estimate of ULCs is made using the empirical relationship between the average ionic charge (*Z*_*av*._) and the solvent-exposed surface area of the corresponding species in solution (*S*):

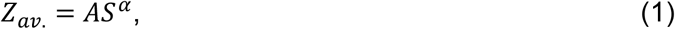

where A is a constant and the empirical value of α is 0.69±0.02.^2^ Using a quasi-spherical approximation for the particle,^1^ the mass (*m*) of the ULC particle can be expressed using its average radius of gyration *R* and effective density *ρ*:

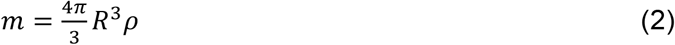

The surface area of the particle can then be expressed as:

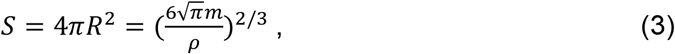

and substituting (3) in (1) allows the average ionic charge to be expressed as:

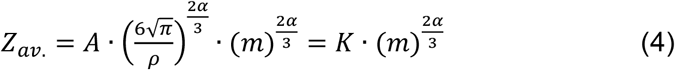

This allows the relationship of the particle mass *m* and the *m/z* value of the corresponding ion to expressed as:

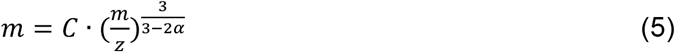

Since in our approximation this *m* vs. *m/z* relationship should hold true for both ULC and SC particles, the mass of the former can be estimated as:

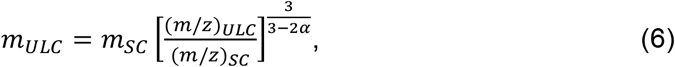

where the empirical value of α is 0.69±0.02,^2^ and all other parameters on the right-hand side of the expression are measured experimentally, providing the following estimate the average ULC mass: *m*_*ULC*_ = 3.0±1.5 MDa.

**Figure S4.**
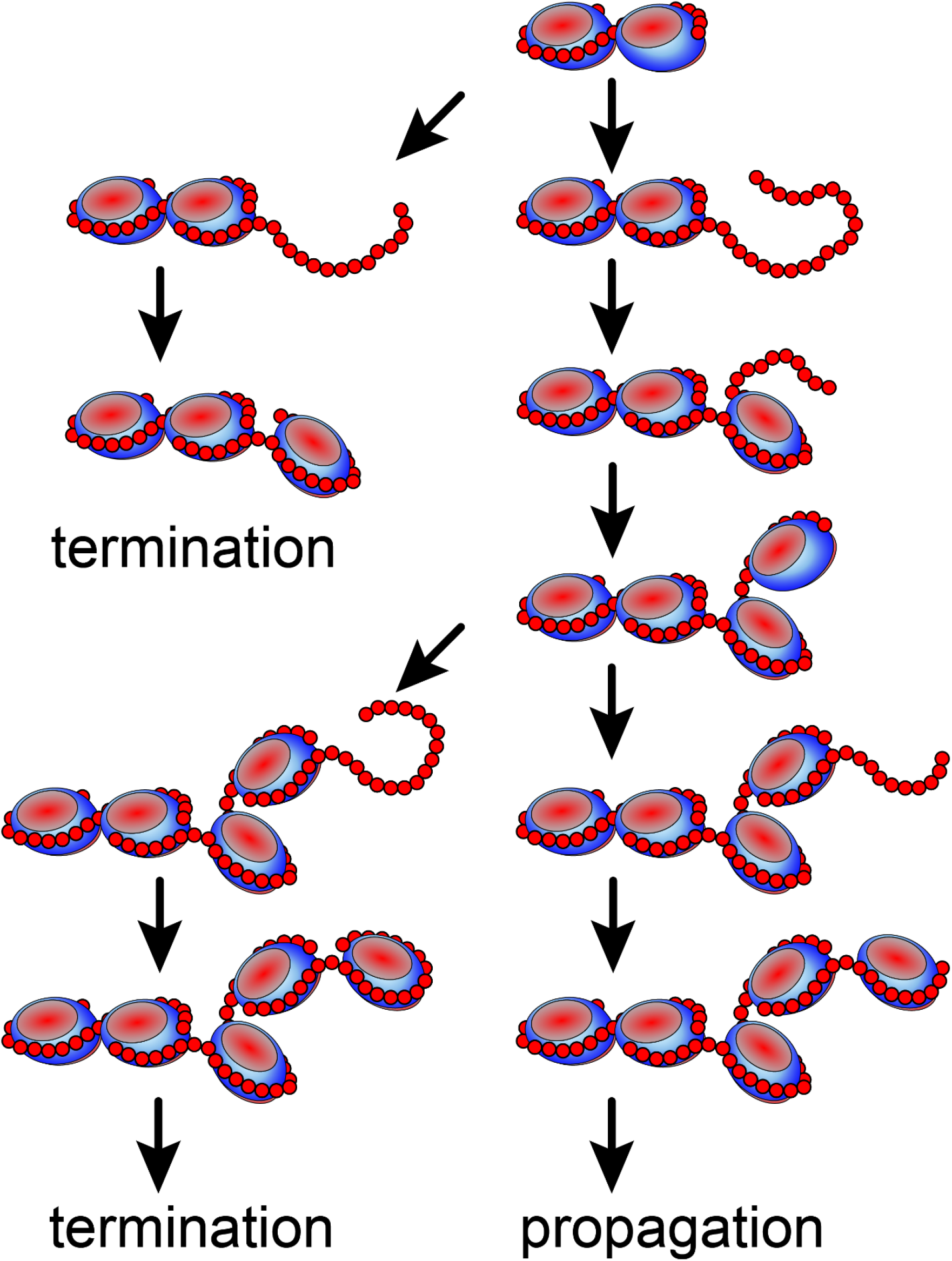
The proposed mechanism of ULC assembly. The successful chain propagation requires that at every step of the assembly process there be a significant mismatch between the “vacant” heparin segment length and the circumference of the PF4 tetramer.

## Notes

### Competing Interest Statement

The authors have declared no competing interest.

